# Molecular dissection of pro-fibrotic signaling identifies the mechanism underlying IL11-driven fibrosis gene translation, reveals non-specific effects of STAT3 and suggests a new mechanism of action for nintedanib

**DOI:** 10.1101/2021.06.10.447846

**Authors:** Anissa A. Widjaja, Sivakumar Viswanathan, Dong Jinrui, Brijesh K. Singh, Jessie Tan, Joyce Goh Wei Ting, David Lamb, Shamini G Shekeran, Benjamin L George, Sebastian Schafer, David Carling, Eleonora Adami, Stuart A. Cook

## Abstract

In fibroblasts, TGFβ1 stimulates *IL11* upregulation that leads to an autocrine loop of IL11-dependent pro-fibrotic protein translation. The signalling pathways downstream of IL11 are contentious and both STAT3 and ERK have been implicated. Here we show that TGFβ1- or IL11-induced ERK activation is consistently associated with fibrogenesis whereas STAT3 phosphorylation (pSTAT3) is unrelated to fibroblast activation. Surprisingly, recombinant human IL11, which has been used extensively in mouse experiments to infer STAT3 activity downstream of IL11, non-specifically increases pSTAT3 in *Il11ra1* null mouse fibroblasts. Pharmacologic inhibition of STAT3 prevents TGFβ1-induced fibrogenesis but this effect was found to reflect fibroblast dysfunction due to severe proteotoxic ER stress. In contrast, inhibition of MEK/ERK prevented fibrosis in the absence of ER stress. TGFβ1-stimulated ERK/mTOR/P70RSK-driven protein translation was IL11-dependent and selectivity for pro-fibrotic protein synthesis was ascribed to an EPRS-related mechanism. In TGFβ1-stimulated fibroblasts, the anti-fibrotic drug nintedanib caused dose-dependent ER stress, reduced pSTAT/pERK and inhibited pro-fibrotic protein translation, similarly to generic STAT3 inhibitors or ER stressors. Pirfenidone, while anti-fibrotic, had no effect on ER stress whereas anti-IL11 inhibited the ERK/mTOR axis while reducing ER stress. These studies discount a specific role for STAT3 in pro-fibrotic signaling, suggest a novel mechanism of action for nintedanib and prioritise further the IL11 pathway as a therapeutic target for fibrosis.

## Introduction

TGFβ1 is one of the most studied human genes and has long been regarded as the dominant fibrogenic factor (Dolgin, 2017). While canonical TGFβ1-driven SMAD activation is central to its activity in fibroblasts, increased protein translation through non-canonical pathways is also important (Chaudhury et al., 2010; Schwarz, 2015; Zhang, 2017). TGFβ1-driven ERK activation has long been recognised as important and more recently STAT3 has been proposed as “*a key integrator of profibrotic signalling*” (Chakraborty et al., 2017; Dees et al., 2012; McHugh, 2017; Zhang, 2017). In 2017, we showed that autocrine IL11 activity is required downstream of TGFβ1-stimulated SMAD activation for fibroblast-to-myofibroblast transformation (Schafer et al., 2017). Intriguingly, the fibrogenic effects of IL11 across stromal cell types are evident only at the translational level (Cook and Schafer, 2020; Lim et al., 2020; Widjaja et al., 2019).

IL11 is a little studied and rather misunderstood member of the IL6 family of proteins (Cook and Schafer, 2020; Widjaja et al., 2020). To signal, IL6 family cytokines bind to their cognate alpha receptors and the receptor:ligand complexes then bind to the common gp130 receptor. Canonical gp130 signaling involves its auto-phosphorylation, recruitment of Janus kinases, STAT3 activation and STAT3 target gene transcription. It thus follows that STAT3 activity could underlie the pro-fibrotic effects of IL11:IL11RA:gp130 signaling in fibroblasts. However, this notion is incongruent with the fact that IL11 does not regulate gene transcription in stromal cells (Cook and Schafer, 2020; Lim et al., 2020; Widjaja et al., 2019).

Here, using neutralizing antibodies, studies of fibroblasts from *Il11ra1* mice, and pharmacological approaches we disentangle fibrogenic effects from non-specific signaling events downstream of TGFβ1 or IL11 and identify the mechanisms underlying IL11-driven pro-fibrotic gene translation. We then apply our findings to better understand the anti-fibrotic activities of pirfenidone and nintedanib, drugs approved for the treatment of fibrotic human diseases whose mechanisms of action (MOA) remain unclear (Roth et al., 2015; Schaefer et al., 2011).

## Results

### Fibrogenesis is associated with IL11-induced ERK but not STAT3 activation

We profiled primary human cardiac fibroblasts (HCFs) stimulated with TGFβ1 over a 24 h period and observed rapid and sustained phosphorylation of SMAD2, bi-phasic activation of ERK (pERK), and late activation of STAT3 (pSTAT3)(**Fig 1A and S1A**). HCFs progressively secreted IL11 over a time course following TGFβ1 stimulation, with marked upregulation by 24h (**Fig S1B**). Immunofluorescence (IF) staining revealed coexpression of IL11RA and gp130 in early passage of HCFs, consistent with autocrine IL11 signaling in HCFs (**Fig 1B**).

**Figure 1.**
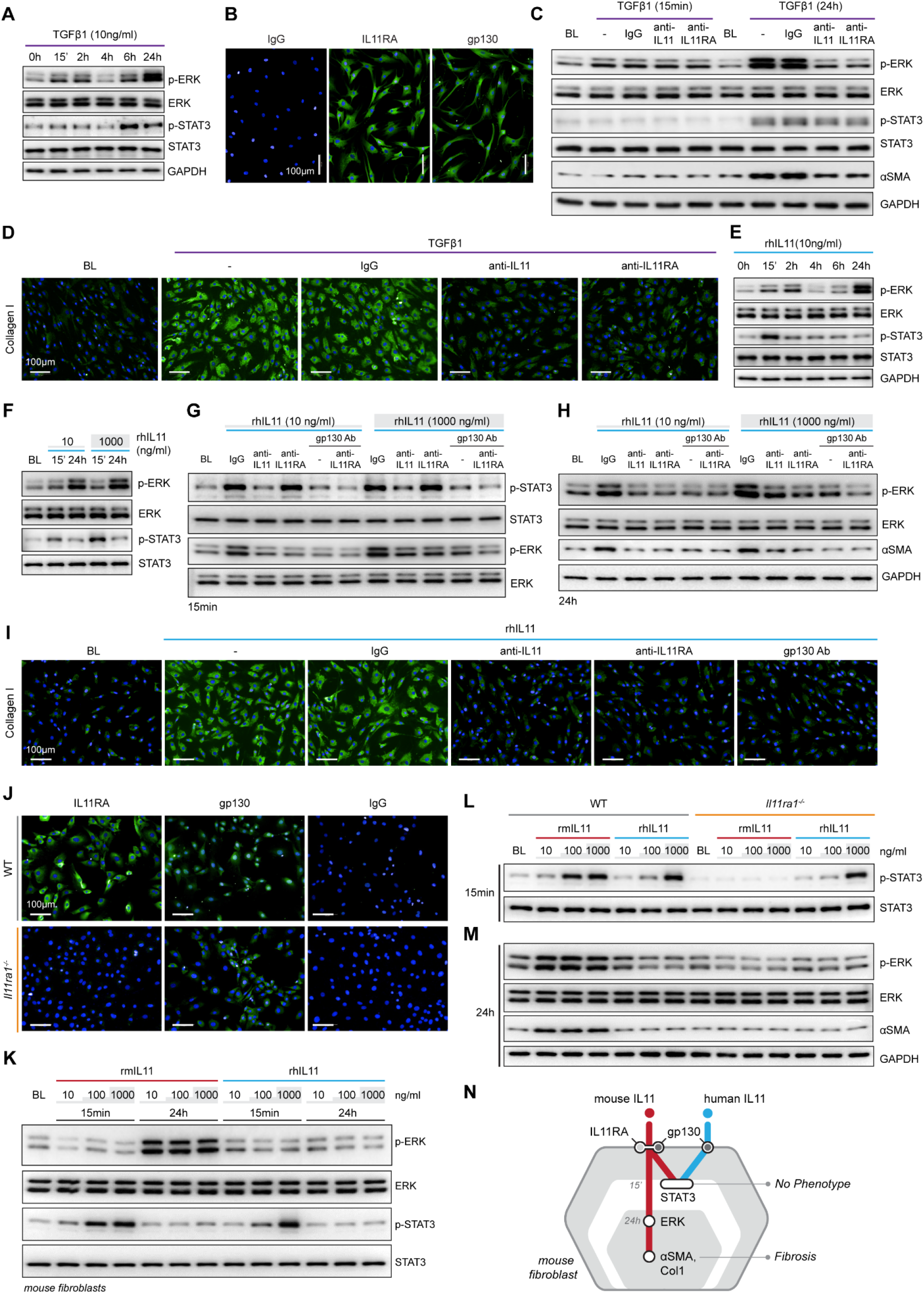
TGFβ1-/IL11-driven fibrogenesis is coincident with activation of ERK, unrelated to STAT3 phosphorylation and shows species-specific effects. (**A**) Western blots of ERK and STAT3 activation in TGFβ1-stimulated HCFs over a time course. (**B**) Representative immunofluorescence (IF) images of IL11RA, and gp130 expression in HCFs (scale bars, 100 μm). (**C-D**) (C) Western blot analysis of p-ERK, ERK, p-STAT3, STAT3, and ⍺SMA and (D) IF images of Collagen I staining in TGFβ1-stimulated HCFs in the presence of either IgG, anti-IL11, or anti-IL11RA. (**E**) Western blots of ERK and STAT3 activation status in IL11-stimulated HCFs over a time course. (**F**) Western blots showing ERK and STAT3 activation status in HCFs following stimulation with low and high dose IL11. (**G-I**) Effects of anti-IL11, anti-IL11RA, or anti-gp130 on inhibiting (G-H) ERK and STAT3 activation and (I) Collagen I Induction in IL11-stimulated HCFs. (**J**) IF images (scale bars, 100 μm) of IL11RA and gp130 in MCFs isolated from wild-type (WT) and *Il11ra1*^-/-^ mice. (**K-M**) Dose-dependent effects of rmIL11 and rhIL11 on (K) ERK and STAT3 activation status in wild-type (WT) MCFs, on (L) STAT3 activation (15 m) and (M) ERK activation and ⍺SMA expression (24h) in WT and *Il11ra1*^-/-^ MCFs. (**N**) Schematic showing the effects of rmIL11 or rhIL11 on signaling and fibrosis in mouse fibroblasts. (A-I) primary HCFs; 24h, (J-M) primary MCFs, (A-M) IL11/TGFβ1 (10 ng/ml), unless otherwise specified. BL: Baseline.

To study the functional relevance of TGFβ1- or IL11-induced ERK and/or STAT3 activation, we stimulated HCFs with TGFβ1 in the presence of neutralizing IL11 or IL11RA antibodies that were specifically developed to inhibit *in vitro* fibrosis phenotypes, agnostic of the underlying pathways (Widjaja et al., 2021, 2019). In TGFβ1-stimulated HCFs, inhibition of IL11 signalling using either anti-IL11 or anti-IL11RA prevented ERK phosphorylation (at the 24h time point) but had no effect on STAT3 activity (**Fig 1C**). Downregulation of ERK activity by anti-IL11 or anti-IL11RA was coincident with a reduction in TGFβ1-stimulated fibroblast-to myofibroblast transformation, as evident from lesser ⍺SMA and Collagen expression (**Fig 1C-D and S1C-E**).

IL11-stimulated HCFs also exhibited ERK activation, which mirrored the effect seen with TGFβ1 (**Fig 1E**). IL11 activated STAT3 in HCFs but its phosphorylation pattern differed from TGFβ1 with an early (15m) increase followed by progressively lower levels. We then stimulated HCFs with a physiological (10ng/ml) or supra-physiological (1000 ng/ml) dose of IL11. ERK was maximally activated with physiological IL11 levels, whereas STAT3 phosphorylation was most pronounced at very high IL11 concentration, questioning the specificity of IL11-stimulated pSTAT3 (**Fig 1F**).

We next studied the effects of anti-IL11 or anti-IL11RA as compared to a commercial anti-IL11 (MAB218), which inhibits cardiac fibroblast-to-myofibroblast transformation (Schafer et al., 2017), as well as a gp130-neutralizing clone (**Fig S1F**). IL11-induced ERK phosphorylation was similarly inhibited by all four neutralizing antibodies (**Fig 1G-H and S1G**). In contrast, MAB218 or anti-IL11RA reduced only ERK phosphorylation but did not prevent STAT3 phosphorylation (**Fig 1G-H and S1G-H**). In IL11 stimulated HCFs, anti-IL11 (two separate clones), anti-IL11RA, and anti-gp130 equally inhibited fibrogenesis as compared to control, as evident from ⍺SMA and Collagen expression levels (**Fig 1H-I and S1I-K**).

Taken together these data confirm and extend the evidence showing that TGFβ1-induced fibrogenesis is IL11:IL11RA:gp130-dependent. They also show that TGFβ1/IL11-stimulated ERK activation is consistently associated with fibrogenesis, whereas STAT3 phosphorylation is not.

### Non-specific effects of high concentration human IL11 in mouse fibroblasts

There are discrepancies in the literature relating to the downstream mediators of IL11, which may relate to the use of recombinant human IL11 (rhIL11) in mouse experiments (Cook and Schafer, 2020). We stimulated primary mouse cardiac fibroblasts (MCFs), which coexpress IL11RA1 and gp130 (**Fig 1J**), with either rhIL11 or recombinant mouse IL11 (rmIL11) over a dose range (ng/ml: 10, 100, 1000) for 15m or 24h. In MCFs, species-matched rmIL11 activated ERK activation with maximal phosphorylation observed at the lowest concentration tested (10ng/ml). rmIL11 also induced STAT3 activation with most notable effects at very high IL11 concentration (1000ng/ml) (**Fig 1K**). In contrast, species-unrelated rhIL11 did not activate ERK in MCFs but instead induced STAT3 phosphorylation when used at high concentrations (≥100ng/ml) (**Fig 1K**).

We studied the species-specific effects in more detail using rhIL11 or rmIL11 on MCFs isolated from *Il11ra1* null mice (*Il11ra1^-/-^*) or wild-type (WT) controls. rmIL11 resulted in STAT3 phosphorylation in WT MCFs at higher doses (>100ng/ml) but did not activate STAT3 in *Il11ra1^-/-^* cells (**Fig 1L**). In WT cells, rmIL11 induced ERK activation and ⍺SMA upregulation at low concentration (10ng/ml), but had no effect on pERK, pSTAT3 of fibrogenesis in *Il11ra1^-/-^* MCFs (**Fig 1M**). In contrast, rhIL11 activated STAT3 in both *Il11ra1^-/-^* and WT fibroblasts when used at high concentrations but had no effect on ERK activation (**Fig 1L-M**). Irrespective of its effects on pSTAT3, rhIL11 did not stimulate fibrosis in MCFs of any genotype.

Overall, these data show unexpected effects of species-unrelated rhIL11 in MCFs. While rhIL11 does not induce pERK or stimulate fibrogenesis in MCFs, high concentrations activate STAT3 in both WT and *Il11ra1^-/-^* fibroblasts. This suggests direct, non-specific binding of high dose rhIL11 to mouse gp130, independent of IL11RA1 (**Fig 1N**).

### Inhibition of STAT3 activity results in fibrogenesis-related ER stress causing fibroblast dysfunction and death

To dissect matters further, we examined the effects of pharmacologic inhibition of either ERK (U0126) or STAT3 (S3I-201) in fibroblasts stimulated with IL11. Surprisingly, inhibition of STAT3 was equally effective in preventing fibrogenesis as ERK inhibition (**Fig 2A-D**). This was unexpected given our earlier findings (**Fig 1**) but consistent with the literature (Chakraborty et al., 2017; Dees et al., 2012). However, while U0126 inhibited IL11-induced ERK but not STAT3 phosphorylation, S3I-201 inhibited both STAT3 and ERK activation, which should not occur with selective inhibition (**Fig 2E**).

**Figure 2.**
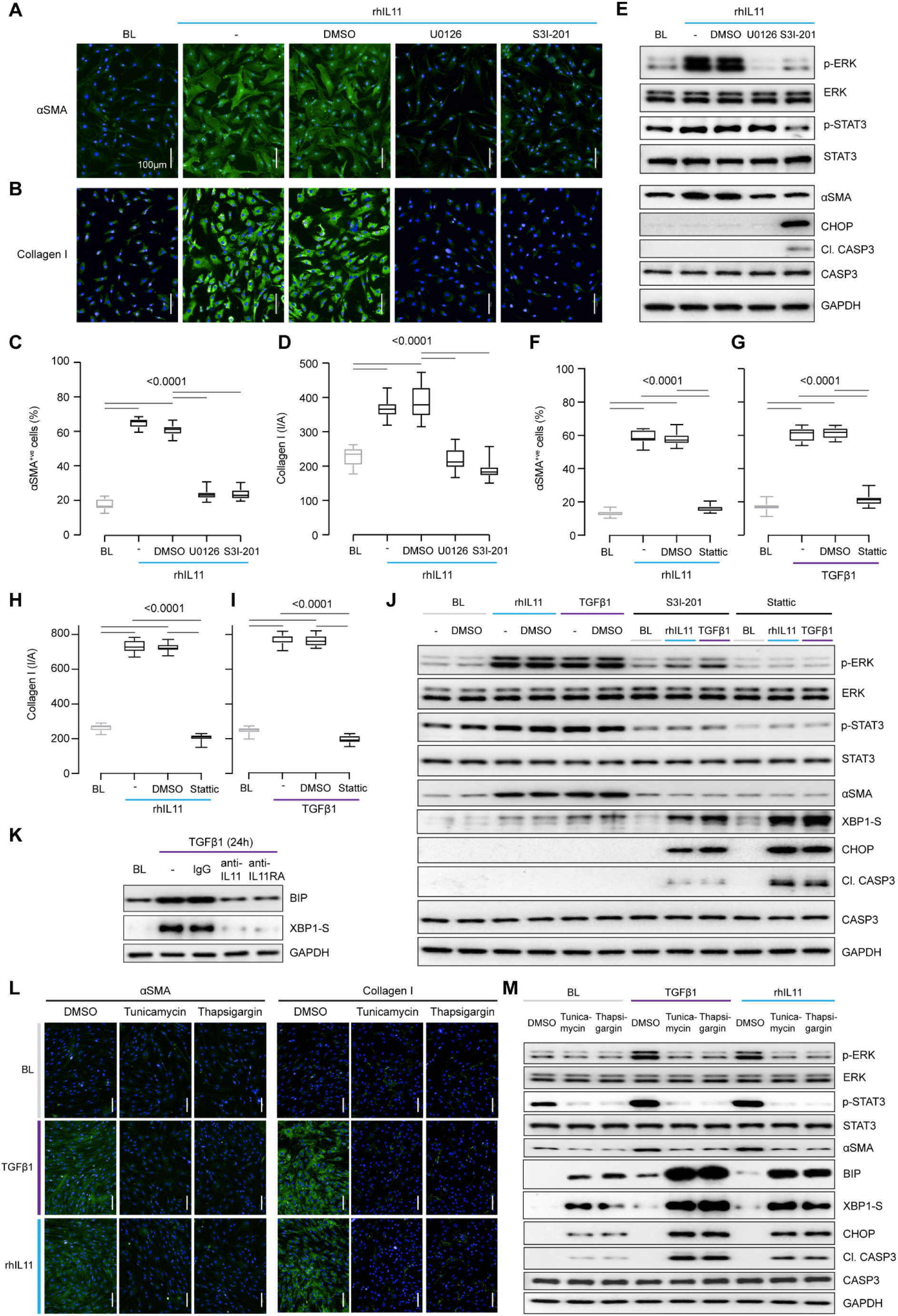
Inhibition of STAT3 in TGFβ1- or IL11-stimulated fibroblasts causes ER stress-related fibroblast dysfunction mimicking effects of generic ER stressors. (**A-E**) Effects of U0126 or S3I-201 on rhIL11-stimulated HCFs. (A-B) IF images (scale bars, 100 μm) and (C-D) quantification of ⍺SMA^+ve^ cells and Collagen I immunostaining. (E) Western blots showing ERK, STAT3, and Caspase3 activation status and ⍺SMA and CHOP protein expression. (**F-I**) Quantification of (F and G) ⍺SMA^+ve^ cells and (H and I) Collagen I immunostaining following stimulation with (F and H) rhIL11 or (G and I) TGFβ1 in the presence of Stattic. (**J**) Effects of S3I-201 or Stattic on the activation of ERK, STAT3, and Caspase3 and on the expression of ⍺SMA, CHOP, and XBP1-S at baseline and in TGFFβ1- or IL11-stimulated HCFs. (**K**) Western blots of BIP and XBP1-S from IgG/anti-IL11/anti-IL11RA-treated TGFβ-stimulated HCFs. (**L-M**) (L) Representative IF images (scale bars, 100 μm) of ⍺SMA^+ve^ cells and Collagen I and (M) Western blot analysis of pERK, ERK, pSTAT3, STAT3, ⍺SMA, BIP, XBP1-S, CHOP, Cleaved Caspase3, Caspase3, and GAPDH from TGFβ1- or rhIL11-stimulated HCFs.(A-M) primary HCFs; 24h; rhIL11/TGFβ1 (10 ng/ml), U0126 (10 μM), S3I-201 (20 μM), Stattic (2.5 μM), IgG/anti-IL11/anti-IL11RA (2 μg/ml), Tunicamycin (5 μg/ml), Thapsigargin (300 nM). (C-D, F-I) Data are shown as box-and-whisker with median (middle line), 25th–75th percentiles (box) and min-max percentiles (whiskers); one-way ANOVA with Tukey’s correction. BL: Baseline

STAT3 activity has been associated with reduced endoplasmic reticulum (ER) stress that occurs with proteotoxicity (Song et al., 2020). We thus examined ER stress in IL11 stimulated HCFs and observed that ERK inhibition with U0126 reduced ⍺SMA induction in the absence of proapoptotic ER stress (CHOP induction and Caspase3 cleavage) (**Fig 2E**). In contrast, S3I-201 inhibited both ERK and STAT3, below baseline, and induced proapoptotic ER stress, which was associated with lesser ⍺SMA expression (**Fig 2E**). Informed by dose-finding experiments (**Fig S2A**), we used a second STAT inhibitor (Stattic) (Schust et al., 2006). Stattic, like S3I-201, prevented TGFβ1 or IL11-induced fibrogenesis (**Fig 2F-I and S2B**) and caused cell death (**S2C**).

Fibroblasts stimulated with pro-fibrotic factors synthesise and secrete large amounts of extracellular proteins that causes a degree of ER stress, which is compensated for by specific chaperone proteins (Baek et al., 2012; Maiers et al., 2017). In keeping with this, the adaptive ER stress proteins XBP1-S and BIP were mildly upregulated in TGFβ1 or IL11-stimulated HCFs (**Fig 2J**). However, in the presence of S31-201 or Stattic, TGFβ1 or IL11 resulted in much more severe and pro-apoptotic ER stress (**Fig 2J**).

As anti-IL11 or anti-IL11RA inhibit IL11-dependent pro-fibrotic protein translation, these antibodies should limit proteotoxic ER stress in HCFs stimulated with pro-fibrotic factors. Indeed, we found that anti-IL11 or anti-IL11RA lowered BIP and XBP1-S expression (**Fig 2K**) in TGFβ1-stimulated HCFs, thus lowering proteotoxic ER stress overall (**Fig S2D**).

The relationship between ER stress and fibrogenesis was further studied using the generic ER stress activators (thapsigargin or tunicamycin), which robustly inhibited TGFβ1-induced fibrogenesis (⍺SMA and Collagen I expression) (**Fig 2L-M**). When given alone, thapsigargin or tunicamycin caused ER stress, as evidenced by elevated BIP, XBP1-S, CHOP and Caspase 3 cleavage, which was increased further with ER protein loading (e.g. with collagen) following TGFβ1 or IL11 stimulation. Interestingly, these generic ER stressors specifically reduced pSTAT3 (but not pERK) levels below baseline when used alone or in combination with TGFβ1 or IL11 (**Fig 2M**).

Taken together these data show that inhibition of IL11-dependent ERK signaling in TGFB1 stimulated fibroblasts reduces pro-fibrotic gene translation and ER stress. In contrast, inhibition of pSTAT3 causes severe ER stress resulting in fibroblast dysfunction that non-specifically prevents myofibroblast transformation, as seen with generic ER stressors.

### TGFβ1-stimulated protein synthesis is IL11-/ERK- and mTOR-dependent

We next examined protein synthesis using the OP-Puro Protein Synthesis Assay (**Fig 3A**). IL11 alone was sufficient to induce protein synthesis in HCFs (**Fig 3B**). Moreover, TGFβ1-stimulated protein synthesis was IL11-dependent and addition of either anti-IL11 or anti-IL11RA was equally effective in inhibiting this (**Fig 3C-D**).

**Figure 3.**
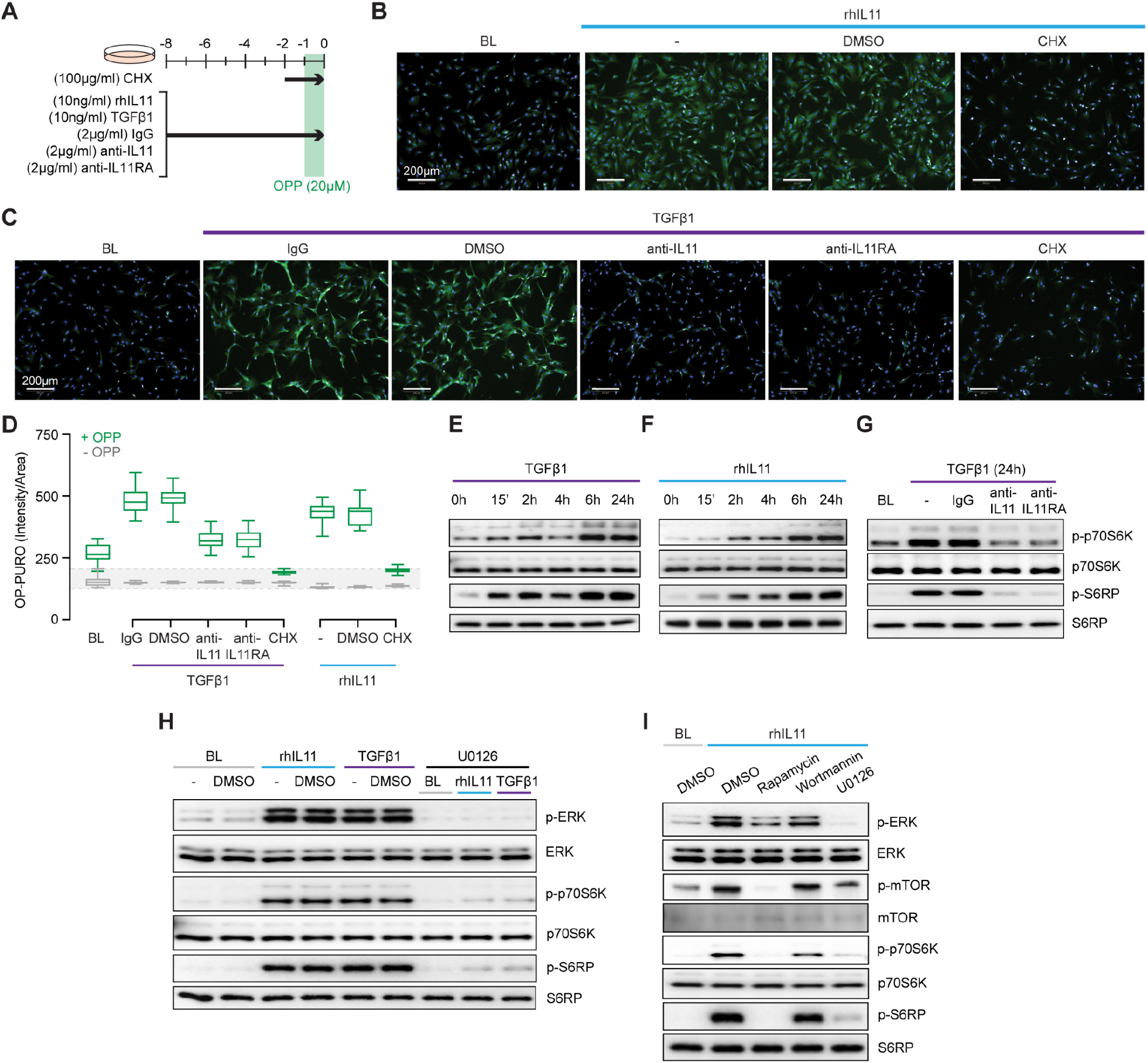
TGFβ1- and IL11-stimulated protein synthesis is ERK- and mTOR-dependent. (**A**) Schematic for experiments shown in B-D. (**B-D**) (B-C) Immunofluorescence images (scale bars, 100 μm) and (D) quantification of Alexa Fluor™ 488 - OPP signal in HCFs following treatments shown in A. (**E-G**) Western blots of p-p70S6K (T389), p70S6K, p-S6RP, S6RP from (E) TGFβ1 or (F) rhIL11-stimulated HCFs over a time course and on (G) IgG, anti-IL11, or anti-IL11RA-treated TGFβ1-stimulated HCFs. (**H**) Effects of U0126 on TGFβ1 or rhIL11-induced ERK, p70S6K (T389), and S6RP activation. (**I**) Comparison effects of Rapamycin, Wortmannin, and U0126 on rhIL11-induced ERK, mTOR, p70S6K, and S6RP activation. (B-I) primary HCFs; IL11/TGFβ1 (10 ng/ml), OPP (20 μM), CHX (100 μg/ml), IgG/anti-IL11/anti-IL11RA (2 μg/ml), U0126 (10 μM), Rapamycin (10 nM), Wortmannin (1μM); (B-D) 8h, (G-I) 24h. (D) Data are shown as box-and-whisker with median (middle line), 25th–75th percentiles (box) and min-max percentiles (whiskers); one-way ANOVA with Tukey’s correction.

Activation of p70S6K is a signalling convergence for canonical protein synthesis pathways that include MEK/ERK, PI3K and AMPK, among others (Wang et al., 2001). Stimulation of HCFs with TGFβ1 resulted in biphasic phosphorylation of p70S6K and of its downstream target S6 ribosomal protein (S6RP) (**Fig 3E**). The effects of IL11 stimulation differed slightly with progressive phosphorylation of p70S6K and S6RP over the time course with maximal activation at 24h (**Fig 3F**). We examined whether TGFβ1-induced p70S6K activation was IL11-dependent, which proved to be the case (**Fig 3G**).

The central importance of ERK activation for TGFβ1- or IL11-induced p70S6K activity was apparent from experiments using the MEK inhibitor U0126 (**Fig 3H**). To explore the pathway components between ERK and p70S6K, we inhibited mTOR with rapamycin and compared its effects with wortmannin. Wortmannin had no effect on p70S6K or S6RP phosphorylation, ruling out a role for PI3K/AKT. In contrast, rapamycin inhibited phosphorylation of mTOR, as expected, and also p70S6K and S6RP activation (**Fig 3I**), similar to effects seen with U0126 (**Fig 3H**). This places mTOR activation downstream of IL11-induced MEK/ERK phosphorylation and upstream of p-p70S6K.

### IL11 stimulates translation of proline-rich fibrosis genes via EPRS

IL11 promotes the translation of pro-fibrotic ECM proteins and of itself but does not non-specifically increase translation of all proteins (Cook and Schafer, 2020; Schafer et al., 2017). Glutamyl-prolyl-tRNA synthetase (EPRS) is important for TGFβ1-stimulated translation of proline-rich proteins, such as collagen, in cardiac fibroblasts (Wu et al., 2020). It is also known that EPRS is phosphorylated by p70S6K that can have non-canonical effects on EPRS function (Arif et al., 2017). This prompted us to examine whether EPRS has a role in IL11-stimulated protein synthesis.

IL11 stimulates its own translation in an autocrine loop and if this were related to EPRS activity then IL11 would need to be proline-rich itself. Examination of human and mouse IL11 revealed a high proline content of both molecules, 11.6% and 8.5 % of amino acids, respectively (**Fig 4A and S3**). Of note, there are four PP and three PPP motifs in human IL11, which are also common in collagen, that cause ribosomal stalling and require EPRS and EIF5A activity for continued translation (Doerfel et al., 2013; Mandal et al., 2014).

**Figure 4.**
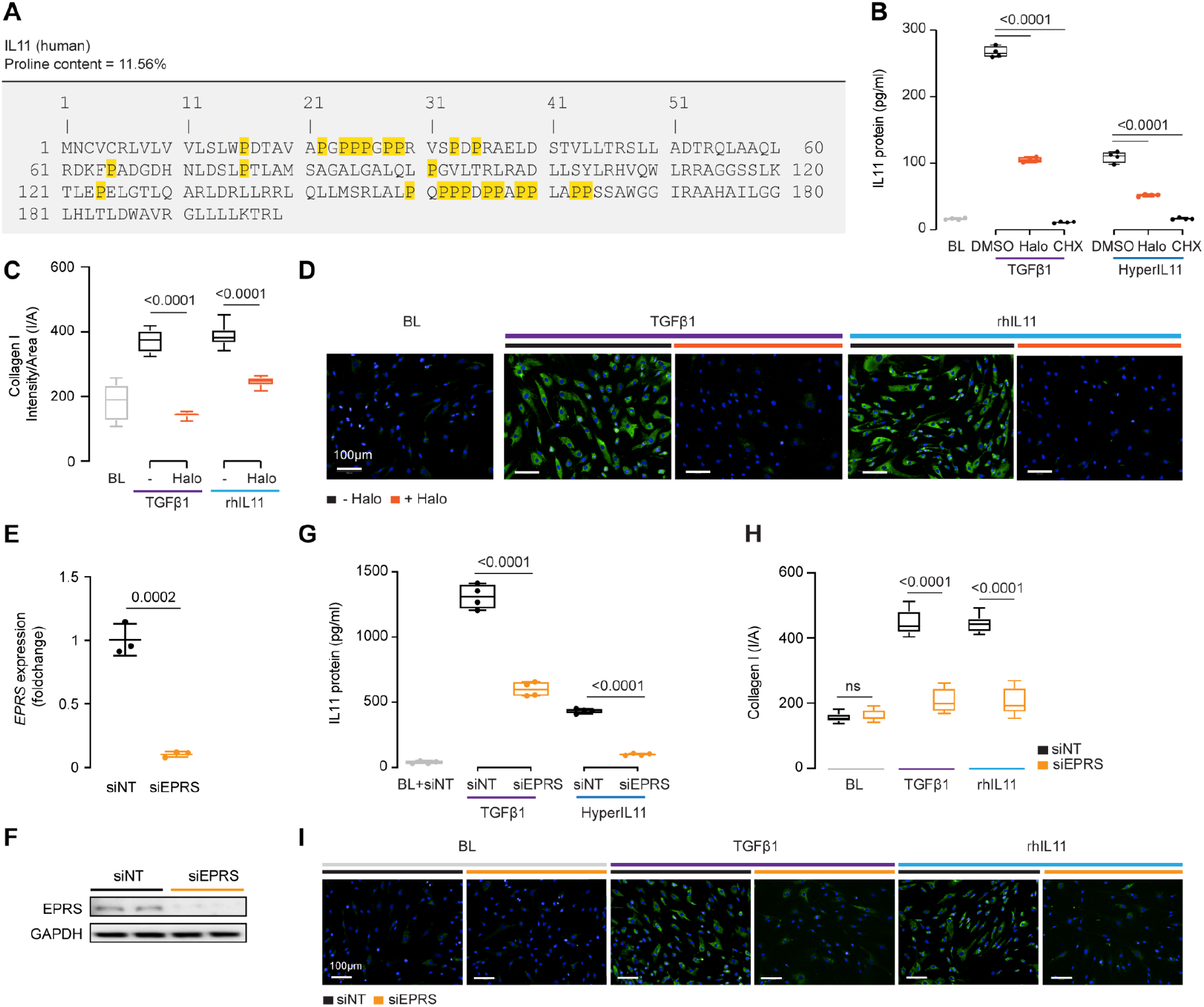
IL11 promotes the translation of proline-rich fibrosis genes, including itself, through EPRS activity. (**A**) Amino acid sequence of human IL11, proline residues highlighted in yellow. (**B**) IL11 levels in the supernatant following TGFβ1 or HyperIL11 stimulation in the presence of Halofuginone (Halo) or Cycloheximide (CHX). (**C-D**) (C) Quantification and (D) immunofluorescence images (scale bars, 100 μm) of Halofuginone-treated TGFβ1/IL11-stimulated HCFs for Collagen I staining. (**E-F**) (E) *EPRS* mRNA and (F) EPRS protein expression levels after knockdown using siEPRS. (**G-I**) (G) IL11 levels in the supernatant, (H) quantification, and (I) immunofluorescence images (scale bars, 100 μm) of Collagen I in stimulated HCFs subjected to siEPRS. (**B-I**) primary HCFs; IL11/TGFβ1/HyperIL11 (10 ng/ml), Cycloheximide (CHX, 100 μg/ml), Halofuginione (100 nM), non-targeting siRNA (siNT)/EPRS siRNA (siEPRS) (12.5 nM); 24h. (B-C, G-H) Data are shown as box-and-whisker with median (middle line), 25th–75th percentiles (box) and min-max percentiles (whiskers); one-way ANOVA with Tukey’s correction; (E) Data are shown as mean ± SD, 2-tailed t-test.

We incubated HCFs with the EPRS inhibitor halofuginone in the presence of TGFβ1 or an IL11:IL11RA fusion construct (HyperIL11), which is not detected by IL11 ELISA (Schafer et al., 2017), and measured IL11 levels in the supernatant. Halofuginone reduced IL11 secretion downstream of either TGFβ1 or HyperIL11 stimulation (**Fig 4B**). Collagen has multiple PPG motifs and we confirmed its induction by TGFβ1 is EPRS-dependent while extending findings to show that IL11-induced collagen secretion also requires EPRS activity (**Fig 4C-D**).

The specificity of halofuginone inhibition of EPRS is established (Keller et al., 2012) but to further confirm our findings we used siRNA. Knockdown of EPRS in HCFs using silencing RNAs against EPRS (siEPRS) was confirmed at the RNA and protein level, as compared to non-targeting siRNA (siNT) (**Fig 4E-F**). As seen with halofuginone, siEPRS reduced TGFβ1- or HyperIL11-induced IL11 secretion and collagen production (**Fig 4G-I**).

Here, we confirmed that collagen synthesis requires EPRS for its translation, identify IL11 as a proline rich molecule containing ribosome stalling motifs and show that EPRS activity underlies IL11 translation.

### Effects of nintedanib, pirfenidone or anti-IL11 on signaling

Pirfenidone and nintedanib are approved drugs for the treatment of fibrotic lung disease but the MOA of these drugs are poorly understood (Roth et al., 2015; Schaefer et al., 2011). Using insights from the experiments described above, we examined the effects of pirfenidone or nintedanib as compared to anti-IL11 in HCFs or lung fibroblasts stimulated with TGFβ1.

In TGFβ1-stimulated HCFs, nintedanib reduced pERK to baseline and pSTAT3 below baseline while inhibiting protein synthesis (p-mTOR, p-p70S6K and pS6RP) and reducing ⍺SMA levels (**Fig 5A**). Unexpectedly, nintedanib caused ER stress as evidenced by increased BIP, XBP1-S, CHOP, and cleaved Caspase 3 levels, similar to that seen with S31-201 or the ER stressors, thapsigargin/tunicamycin (**Fig 2**).

**Figure 5.**
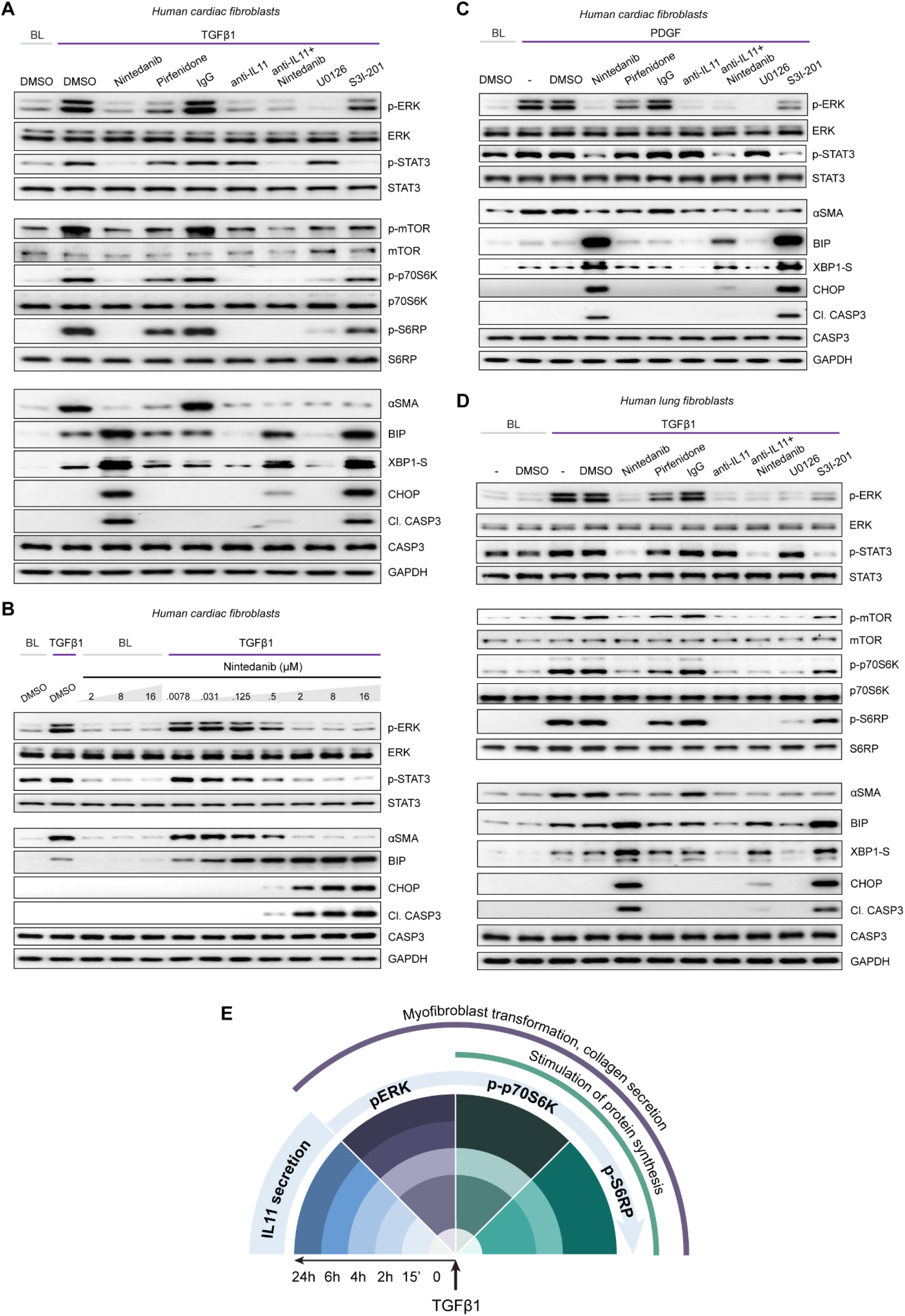
Nintedanib, Pirfenidone and anti-IL11 have different anti-fibrotic mechanisms of action in fibroblasts. (**A&D**) Western blots showing activation status of ERK, STAT3, mTOR, p70S6K (T389), S6RP, and Caspase3, and protein expression of ⍺SMA, BIP, XBP1-S, and CHOP following treatment with nintedanib, pirfenidone, IgG, anti-IL11, a combination of nintedanib, anti-IL11, U0126 or S3I-201 from TGFβ1-stimulated (A) HCFs and (D) human lung fibroblasts. (**B**) Western blots of p-ERK, ERK, p-STAT3, STAT3, BIP, CHOP, Cleaved Caspase 3, Caspase 3, and GAPDH from HCFs treated with different concentrations of nintedanib in the absence or presence of TGFβ1. (**C**) Western blots of p-ERK, ERK, p-STAT3, STAT3, BIP, CHOP, Cleaved Caspase 3, Caspase 3, XBP1-S, and GAPDH from PDGF-stimulated HCFs. (**E**) Schematic overview and timeline of key pro-fibrotic events that occur downstream of TGFβ1-stimulation in human fibroblasts. (A-C) primary HCFs, (D) primary human lung fibroblasts. (A-D) 24 hours; IL11/TGFβ1 (10 ng/ml), IgG/anti-IL11 (2 μg/ml), nintedanib (2 μM), pirfenidone (0.3 mg/ml), U0126 (10 μM), S3I-201 (20 μM), unless otherwise specified.

In contrast, pirfenidone mildly diminished pERK and pSTAT3 and had limited effect on pmTOR, p-p70S6K or pS6RP as compared to nintedanib or anti-IL11. Furthermore, pirfenidone was not associated with increased ER stress over TGFβ1 treatment alone. Anti-IL11 recapitulated earlier findings: inhibiting protein synthesis while lowering TGFβ1-induced proteotoxic ER stress (**Fig 2 and 3**). In addition, anti-IL11 reduced nintedanib-associated ER stress when used in combination (**Fig 5A**).

Nintedanib is not known to cause ER stress and while the concentration we used (2μM) is similar to that commonly applied (Wollin et al., 2015), we probed matters further using a dose-response (nintedanib: 16μM-7.8nM; 4-fold dilutions) (**Fig 5B**). Nintedanib at a concentration up to 16μM had no effect on ER stress in quiescent fibroblasts. However, in TGFβ1-stimulated HCFs, nintedanib began to inhibit ⍺SMA expression at 125nM and complete inhibition was observed at 2μM. Over the same concentration range, BIP was progressively induced and at the higher end of the range (0.5μM and 2μM), pro-apoptotic ER stress (CHOP/cleaved Caspase3) upregulation was more apparent. Nintedanib also inhibited pERK and pSTAT in a dose-dependent manner, which was inversely related to the induction of ER stress markers.

The MOA of Nintedanib is thought through inhibition of PDGF and FGF signaling in fibroblasts. We thus examined the effects of Nintedanib on PDGF-stimulated HCF signaling and observed very similar effects to those seen with TGFβ1 stimulation: lesser pSTAT and pERK as well as induction of pro-apoptotic ER stress, which was associated with lesser ⍺SMA induction (**Fig 5C**).

Experiments were repeated in TGFβ1-stimulated human lung fibroblasts to exclude organ-of-origin-specific effects in fibroblasts (**Fig 5D**). In lung fibroblasts, nintedanib reduced pSTAT3 below baseline, diminished pERK and limited synthesis of ⍺SMA while inducing pro-apoptotic ER stress, which was lesser with co-administration of anti-IL11. Pirfenidone slightly reduced pERK and pSTAT3 and had lesser effects on the protein synthesis pathway than nintedanib or anti-IL11, as seen in HCFs. Pirfenidone reduced ⍺SMA expression, as expected, but had no effect over TGFβ1 on markers of ER stress.

## Discussion

In this study, we set out to identify the signaling pathways by which TGFβ1-induced IL11 activity increases profibrotic gene translation. In particular, we wished to dissect the relative contributions of ERK from STAT3 downstream of IL11. IL11 is a little studied and somewhat misunderstood cytokine (Cook and Schafer, 2020; Widjaja et al., 2020), as exemplified by publications on cardiac and renal fibrosis where it was originally [and erroneously] reported as anti-fibrotic (Obana et al., 2010). We have speculated that confusion relating to IL11 biology in general, and IL11-related STAT3 phosphorylation in particular, may stem from the use of human IL11 in murine cells and models, as noted previously (Cook and Schafer, 2020; Widjaja et al., 2020).

Here we show that species-matched IL11 transiently increases pSTAT3 and TGFβ1 induces late onset STAT3 phosphorylation in fibroblasts, but these events are unrelated to fibrosis. Notably, rhIL11 activates STAT3 in both WT and *I11ra1*-deleted MCFs by binding directly to gp130 but does not activate ERK, which rmIL11 does. This ‘off-target’ effect of rhIL11 on pSTAT3 in mouse cells is exacerbated by the recent finding that rhIL11 paradoxically antagonises endogenous IL11 signaling in mouse models of disease (Widjaja et al., 2021). To confound matters further, IL11RA is rapidly lost from fibroblasts in culture, even at early passage, and very high doses of rhIL11 are often used in mouse experiments.

In phenotypic studies, we were puzzled to observe that STAT3 inhibition prevented fibrogenesis, which was not consistent with our signalling data, but we noted that S31-201 reduced ERK phosphorylation. This could reflect complexities of signal pathway cross-talk or more generalised fibroblast dysfunction as inhibition of STAT3 is linked with pro-apoptotic ER stress (Song et al., 2020). TGFβ1 or IL11 stimulation resulted in mild proteotoxic ER stress, as expected, but this homeostatic mechanism transitioned to pro-apoptotic ER stress with STAT3 inhibition. Tellingly, incubation of HCFs with thapsigargin or tunicamycin resulted in severe ER stress and inhibited fibrogenesis when administered with TGFβ1 or IL11. Hence, while we confirm that inhibition of STAT3 reduces fibrosis in keeping with the literature (Dees et al., 2012; McHugh, 2017), this effect appears ER-stress related and non-specific.

In contrast to the effects of STAT3 inhibitors, anti-IL11 or anti-IL11RA reduced both fibrogenesis and ER stress in TGFβ1-stimulated HCFs. Inhibition of IL11 signalling was associated with lesser pERK but not pSTAT3 and mimicked the antifibrotic effects seen with the MEK/ERK inhibitor, U0126. We confirmed that IL11 is centrally important for protein translation in TGFβ1-stimulated HCFs and showed that activation of mTOR, P70S6K and S6RP is both IL11-dependent and ERK-driven. Our data identify IL11-stimulated ERK-dependent activation of mTOR as a major mechanism for the translational effects of IL11 (**Fig 5E**). Whether ERK activates mTOR directly or there is greater upstream complexity, remains to be elucidated (Du et al., 2008).

Protein synthesis is the most energy-intensive process in growing cells and thus performed in a selective manner (Buttgereit and Brand, 1995). In fibroblasts, EPRS, the bifunctional glutamate/proline tRNA ligase, specifically controls translation of proline rich ECM genes, such as collagen (Wu et al., 2020). We found that IL11 has an unusually high proline content and encodes motifs associated with ribosome stalling that require EPRS and eIF5A for efficient translation (Doerfel et al., 2013). As with collagen, we found that IL11 requires EPRS, a P70S6K target (Arif et al., 2017), for its translation and secretion, which is thus coordinated with that of ECM proteins during fibrogenesis.

Nintedanib was first developed as an anticancer drug and pirfenidone for inflammation and it was only later that these drugs were found to have anti-fibrotic effects (Roth et al., 2015; Schaefer et al., 2011). Nintedanib is a receptor tyrosine kinase inhibitor (TKI), with activity against VEGFR, PDGFR and FGFR with IC50 values of 13-34nM, 59-65nM and 37-108nM, respectively (Hilberg et al., 2008). Nevertheless, in a panel of 33 additional kinases, nintedanib additionally inhibits Flt-3, Lck, Lyn and Src with IC50 values of 26, 16, 196 and 156nM respectively (Hilberg et al., 2008). Whereas pirfenidone is thought to have antioxidant activity. The anti-fibrotic MOA of both drugs remains an issue of much debate.

In TGFβ1- or PDGF-stimulated fibroblasts, nintedanib reduced pSTAT3 below baseline, which was accompanied by induction of pro-apoptotic ER stress. These features were similar to those seen with the use of either STAT3 inhibitors or ER stressors. Interestingly, haloperidol was recently shown to inhibit TGFβ1-induced fibroblast activation and its MOA related to the induction of ER stress (Rehman et al., 2019). The mechanism and directionality by which nintedanib inhibits pSTAT3 and induces ER stress remain to be determined but could relate to TKI activity against, perhaps ER-localised, JAK. We propose that nintedanib’s anti-fibrotic MOA in TGFβ1-stimulated fibroblasts is perhaps due to induction ER stress. However, it should be recognised that nintedanib concentrations equivalent to its C_max_ in patients (~25nM (Wind et al., 2019)) had no effect on fibroblast activation in our assays and a fibroblast-independent MOA may be relevant for therapeutic effect.

Here we dissected the pro-fibrogenic translational-specific signaling activity downstream of IL11 and show the ERK/mTOR/p70S6K axis to be of importance while discounting a role for STAT3. We end by proposing therapeutic inhibition of IL11 as an alternative approach for treating fibrosis that leverages a MOA that is differentiated from those of approved anti-fibrotic drugs. Given the safety data evident from humans and mice with *IL11RA* or *IL11* loss of function and the lack of toxicities with long term anti-IL11 administration (Widjaja et al., 2020, 2019), it is hoped that IL11-targeting approaches may have lesser side effects than current therapies.

## Data availability

All data are provided in the main manuscript or as supporting information.

## Acknowledgement

This research was supported by the National Medical Research Council (NMRC), Singapore STaR awards (NMRC/STaR/0029/2017), NMRC Centre Grant to the NHCS, MOH‐CIRG18nov‐0002, MRC-LMS (UK), Goh Foundation, Tanoto Foundation to S.A.C. A.A.W. is supported by NMRC/OFYIRG/0053/2017.

## References

Arif A, Terenzi F, Potdar AA, Jia J, Sacks J, China A, Halawani D, Vasu K, Li X, Brown JM, Chen J, Kozma SC, Thomas G, Fox PL. 2017. EPRS is a critical mTORC1-S6K1 effector that influences adiposity in mice. Nature 542:357–361.

Baek HA, Kim DS, Park HS, Jang KY, Kang MJ, Lee DG, Moon WS, Chae HJ, Chung MJ. 2012. Involvement of endoplasmic reticulum stress in myofibroblastic differentiation of lung fibroblasts. Am J Respir Cell Mol Biol 46:731–739.

Buttgereit F, Brand MD. 1995. A hierarchy of ATP-consuming processes in mammalian cells. Biochem J 312 (Pt 1):163–167.

Chakraborty D, Šumová B, Mallano T, Chen C-W, Distler A, Bergmann C, Ludolph I, Horch RE, Gelse K, Ramming A, Distler O, Schett G, Šenolt L, Distler JHW. 2017. Activation of STAT3 integrates common profibrotic pathways to promote fibroblast activation and tissue fibrosis. Nat Commun 8:1130.

Chaudhury A, Hussey GS, Ray PS, Jin G, Fox PL, Howe PH. 2010. TGF-beta-mediated phosphorylation of hnRNP E1 induces EMT via transcript-selective translational induction of Dab2 and ILEI. Nat Cell Biol 12:286–293.

Cook SA, Schafer S. 2020. Hiding in Plain Sight: Interleukin-11 Emerges as a Master Regulator of Fibrosis, Tissue Integrity, and Stromal Inflammation. Annu Rev Med 71:263–276.

Dees C, Tomcik M, Palumbo-Zerr K, Distler A, Beyer C, Lang V, Horn A, Zerr P, Zwerina J, Gelse K, Distler O, Schett G, Distler JHW. 2012. JAK-2 as a novel mediator of the profibrotic effects of transforming growth factor β in systemic sclerosis. Arthritis Rheum 64:3006–3015.

Doerfel LK, Wohlgemuth I, Kothe C, Peske F, Urlaub H, Rodnina MV. 2013. EF-P is essential for rapid synthesis of proteins containing consecutive proline residues. Science 339:85–88.

Dolgin E. 2017. The most popular genes in the human genome. Nature 551:427–431.

Du J, Guan T, Zhang H, Xia Y, Liu F, Zhang Y. 2008. Inhibitory crosstalk between ERK and AMPK in the growth and proliferation of cardiac fibroblasts. Biochem Biophys Res Commun 368:402–407.

Hilberg F, Roth GJ, Krssak M, Kautschitsch S, Sommergruber W, Tontsch-Grunt U, Garin-Chesa P, Bader G, Zoephel A, Quant J, Heckel A, Rettig WJ. 2008. BIBF 1120: triple angiokinase inhibitor with sustained receptor blockade and good antitumor efficacy. Cancer Res 68:4774–4782.

Keller TL, Zocco D, Sundrud MS, Hendrick M, Edenius M, Yum J, Kim Y-J, Lee H-K, Cortese JF, Wirth DF, Dignam JD, Rao A, Yeo C-Y, Mazitschek R, Whitman M. 2012. Halofuginone and other febrifugine derivatives inhibit prolyl-tRNA synthetase. Nat Chem Biol 8:311–317.

Lim W-W, Corden B, Ng B, Vanezis K, D’Agostino G, Widjaja AA, Song W-H, Xie C, Su L, Kwek X-Y, Tee NGZ, Dong J, Ko NSJ, Wang M, Pua CJ, Jamal MH, Soh B, Viswanathan S, Schafer S, Cook SA. 2020. Interleukin-11 is important for vascular smooth muscle phenotypic switching and aortic inflammation, fibrosis and remodeling in mouse models. Sci Rep 10:17853.

Maiers JL, Kostallari E, Mushref M, deAssuncao TM, Li H, Jalan-Sakrikar N, Huebert RC, Cao S, Malhi H, Shah VH. 2017. The unfolded protein response mediates fibrogenesis and collagen I secretion through regulating TANGO1 in mice. Hepatology 65:983–998.

Mandal A, Mandal S, Park MH. 2014. Genome-wide analyses and functional classification of proline repeat-rich proteins: potential role of eIF5A in eukaryotic evolution. PLoS One 9:e111800.

McHugh J. 2017. Systemic sclerosis: STAT3 - A key integrator of profibrotic signalling. Nat Rev Rheumatol.

Obana M, Maeda M, Takeda K, Hayama A, Mohri T, Yamashita T, Nakaoka Y, Komuro I, Takeda K, Matsumiya G, Azuma J, Fujio Y. 2010. Therapeutic activation of signal transducer and activator of transcription 3 by interleukin-11 ameliorates cardiac fibrosis after myocardial infarction. Circulation 121:684–691.

Rehman M, Vodret S, Braga L, Guarnaccia C, Celsi F, Rossetti G, Martinelli V, Battini T, Long C, Vukusic K, Kocijan T, Collesi C, Ring N, Skoko N, Giacca M, Del Sal G, Confalonieri M, Raspa M, Marcello A, Myers MP, Crovella S, Carloni P, Zacchigna S. 2019. High-throughput screening discovers antifibrotic properties of haloperidol by hindering myofibroblast activation. JCI Insight 4. doi:10.1172/jci.insight.123987

Roth GJ, Binder R, Colbatzky F, Dallinger C, Schlenker-Herceg R, Hilberg F, Wollin S-L, Kaiser R. 2015. Nintedanib: from discovery to the clinic. J Med Chem 58:1053–1063.

Schaefer CJ, Ruhrmund DW, Pan L, Seiwert SD, Kossen K. 2011. Antifibrotic activities of pirfenidone in animal models. Eur Respir Rev 20:85–97.

Schafer S, Viswanathan S, Widjaja AA, Lim W-W, Moreno-Moral A, DeLaughter DM, Ng B, Patone G, Chow K, Khin E, Tan J, Chothani SP, Ye L, Rackham OJL, Ko NSJ, Sahib NE, Pua CJ, Zhen NTG, Xie C, Wang M, Maatz H, Lim S, Saar K, Blachut S, Petretto E, Schmidt S, Putoczki T, Guimarães-Camboa N, Wakimoto H, van Heesch S, Sigmundsson K, Lim SL, Soon JL, Chao VTT, Chua YL, Tan TE, Evans SM, Loh YJ, Jamal MH, Ong KK, Chua KC, Ong B-H, Chakaramakkil MJ, Seidman JG, Seidman CE, Hubner N, Sin KYK, Cook SA. 2017. IL-11 is a crucial determinant of cardiovascular fibrosis. Nature 552:110–115.

Schust J, Sperl B, Hollis A, Mayer TU, Berg T. 2006. Stattic: a small-molecule inhibitor of STAT3 activation and dimerization. Chem Biol 13:1235–1242.

Schwarz RI. 2015. Collagen I and the fibroblast: high protein expression requires a new paradigm of post-transcriptional, feedback regulation. Biochem Biophys Rep 3:38–44.

Song M, Wang C, Yang H, Chen Y, Feng X, Li B, Fan H. 2020. P-STAT3 Inhibition Activates Endoplasmic Reticulum Stress-Induced Splenocyte Apoptosis in Chronic Stress. Front Physiol 11:680.

Wang L, Gout I, Proud CG. 2001. Cross-talk between the ERK and p70 S6 kinase (S6K) signaling pathways. MEK-dependent activation of S6K2 in cardiomyocytes. J Biol Chem 276:32670–32677.

Widjaja AA, Chothani SP, Cook SA. 2020. Different roles of interleukin 6 and interleukin 11 in the liver: implications for therapy. Hum Vaccin Immunother 16:2357–2362.

Widjaja AA, Dong J, Adami E, Viswanathan S, Ng B, Pakkiri LS, Chothani SP, Singh BK, Lim WW, Zhou J, Shekeran SG, Tan J, Lim SY, Goh J, Wang M, Holgate R, Hearn A, Felkin LE, Yen PM, Dear JW, Drum CL, Schafer S, Cook SA. 2021. Redefining IL11 as a regeneration-limiting hepatotoxin and therapeutic target in acetaminophen-induced liver injury. Science Translational Medicine. doi:10.1126/scitranslmed.aba8146

Widjaja AA, Singh BK, Adami E, Viswanathan S, Dong J, D’Agostino GA, Ng B, Lim WW, Tan J, Paleja BS, Tripathi M, Lim SY, Shekeran SG, Chothani SP, Rabes A, Sombetzki M, Bruinstroop E, Min LP, Sinha RA, Albani S, Yen PM, Schafer S, Cook SA. 2019. Inhibiting Interleukin 11 Signaling Reduces Hepatocyte Death and Liver Fibrosis, Inflammation, and Steatosis in Mouse Models of Non-Alcoholic Steatohepatitis. Gastroenterology. doi:10.1053/j.gastro.2019.05.002

Wind S, Schmid U, Freiwald M, Marzin K, Lotz R, Ebner T, Stopfer P, Dallinger C. 2019. Clinical Pharmacokinetics and Pharmacodynamics of Nintedanib. Clin Pharmacokinet 58:1131–1147.

Wollin L, Wex E, Pautsch A, Schnapp G, Hostettler KE, Stowasser S, Kolb M. 2015. Mode of action of nintedanib in the treatment of idiopathic pulmonary fibrosis. Eur Respir J 45:1434–1445.

Wu J, Subbaiah KCV, Xie LH, Jiang F, Khor E-S, Mickelsen D, Myers JR, Tang WHW, Yao P. 2020. Glutamyl-Prolyl-tRNA Synthetase Regulates Proline-Rich Pro-Fibrotic Protein Synthesis During Cardiac Fibrosis. Circ Res 127:827–846.

Zhang YE. 2017. Non-Smad Signaling Pathways of the TGF-β Family. Cold Spring Harb Perspect Biol 9. doi:10.1101/cshperspect.a022129

