## Supplementary File for "Molecular dissection of pro-fibrotic signaling identifies the mechanism underlying IL11-driven fibrosis gene translation, reveals non-specific effects of STAT3 and suggests a new mechanism of action for nintedanib"

### Supplementary Materials

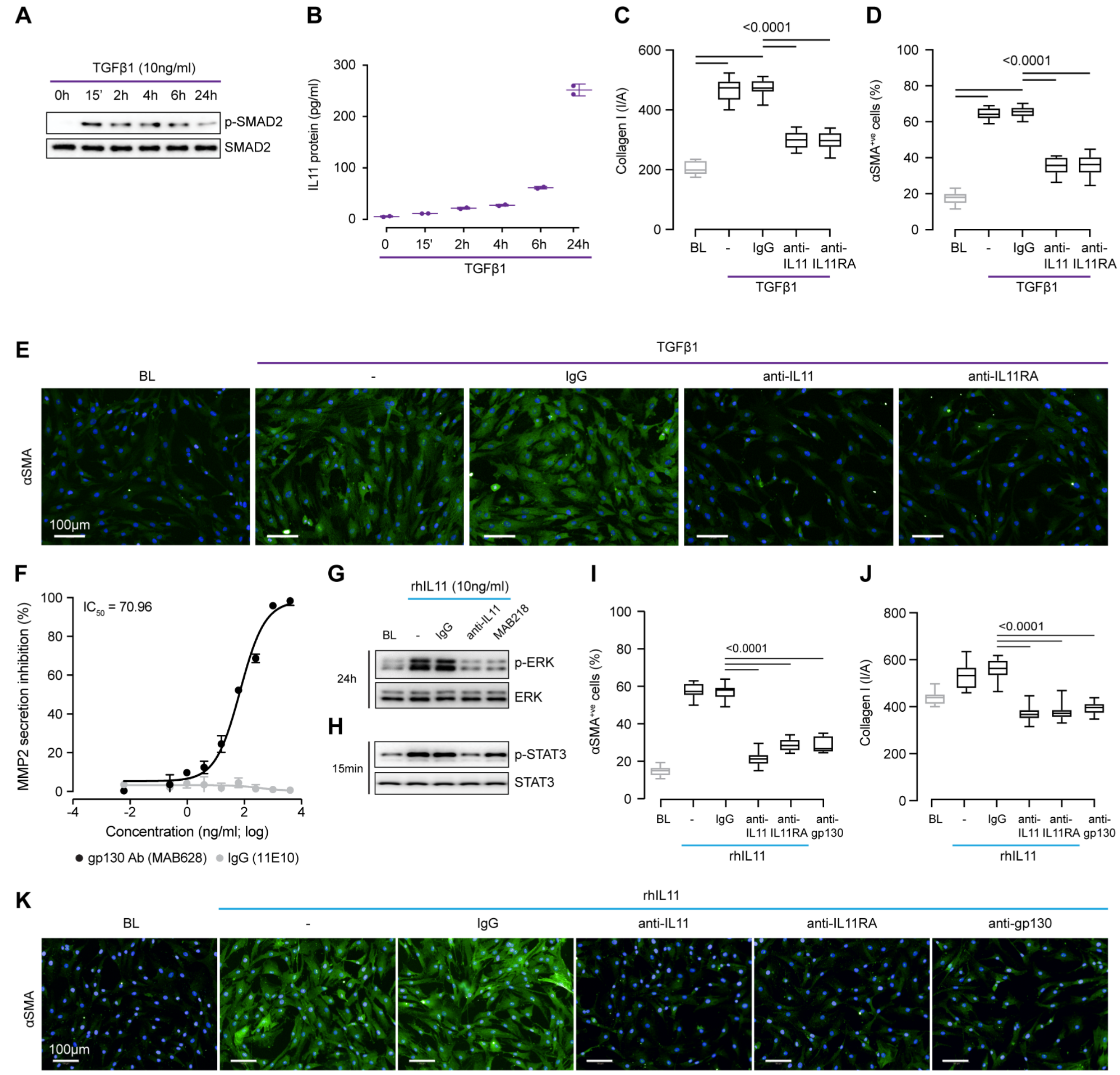


**Figure S1 TGFβ1 stimulates myofibroblast transformation in an IL11- and ERK- dependent manner.** (**A-B**) (A) SMAD2 activation status in the lysates and (B) levels of IL11 protein in the supernatant from TGFβ1-stimulated HCFs over a time course. (**C-E**) Quantification of (C) Collagen I staining and (D) $\alpha$SMA$^{+ve}$ cells and (E) representative IF images of $\alpha$SMA staining in TGFβ1-stimulated HCFs in the presence of either IgG, anti-IL11, or anti-IL11RA. (**F**) Dose-response curve and $IC_{50}$ value of IgG and anti-human gp130 (MAB628, R&D Systems, range: 61 pg ml$^{-1}$ to 4 μg ml$^{-1}$; 4-fold dilution) in inhibiting MMP2 secretion in the supernatant from rmIL11-stimulated primary HCFs. (**G-H**) Western blot analysis of (G) ERK (24h) and (G) STAT3(15m) activation status in IL11-stimulated HCFs in the presence of IgG, anti-IL11, or a commercial neutralizing anti-IL11 (MAB218).(**I-K**) Quantification of the effects of anti-IL11, anti-IL11RA, or anti-gp130 on (I) $\alpha$SMA and (J) Collagen I Induction in IL11-stimulated HCFs and (K) their respective representative immunofluorescence images. (A-K) primary HCFs; 24h; IL11/TGFβ1 (10 ng/ml), IgG/anti-IL11/anti-IL11RA/MAB218/anti-gp130 (2 μg/ml), unless otherwise specified. (B, F) Data are shown as mean ± SD, (C-D, I-J) data are shown as box-and-whisker with median (middle line), 25th–75th percentiles (box) and min-max percentiles (whiskers); one-way ANOVA with Tukey's correction. BL: baseline

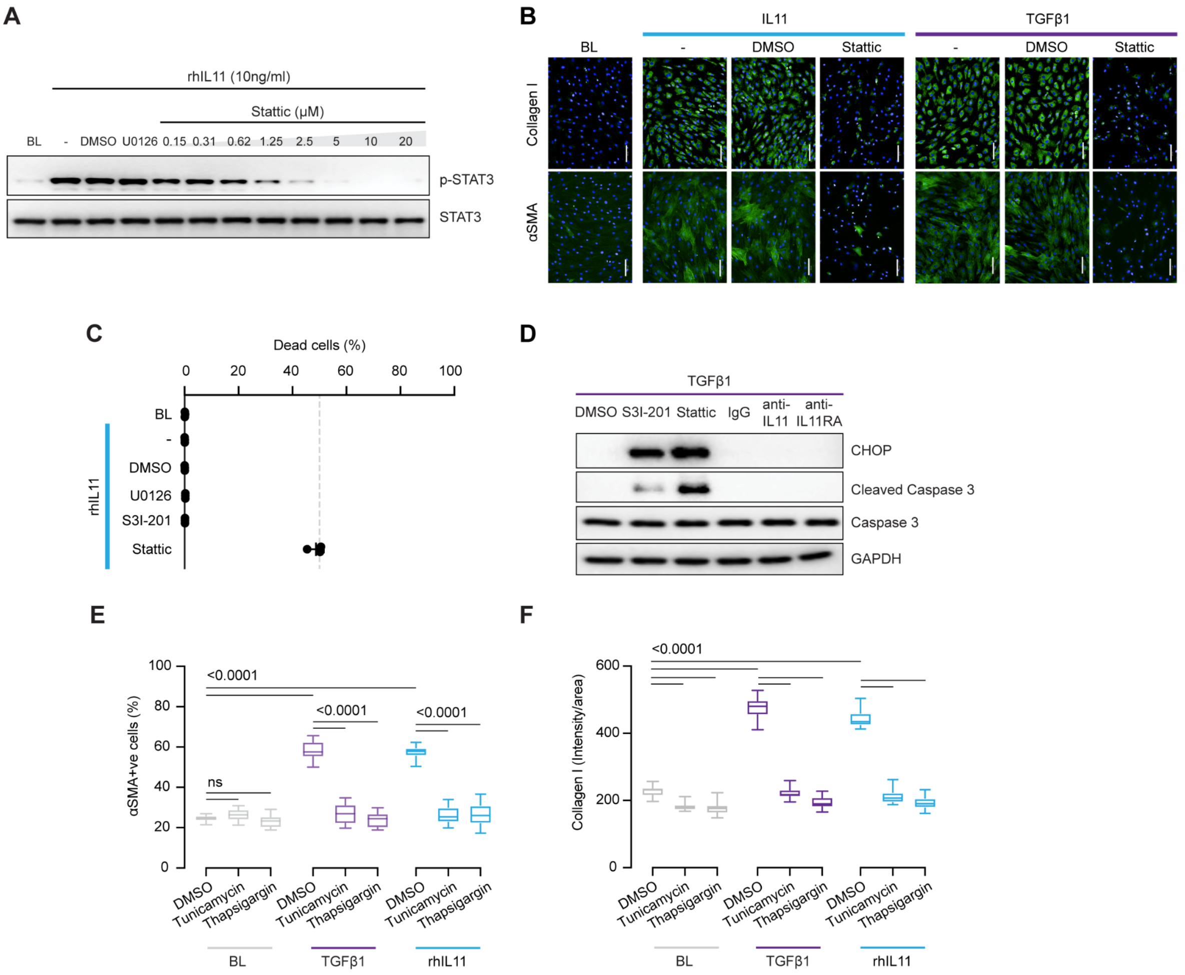


**Figure S2 STAT3 inhibition causes ER stress-related fibroblast dysfunction and cell death.** (**A**) Dose-dependent effects of increasing concentration of Stattic on STAT3 phosphorylation in IL11-stimulated HCFs at 15m time point. (**B**) Representative fluorescence images of αSMA and Collagen I immunostaining in HCFs following stimulation with rhIL11 or TGFβ1 in the presence of Stattic. (**C**) Effects of U0126, S3I-201, or Stattic on cell viability as assayed by live/dead cell staining; data are shown as mean ± SD. (**D**) Comparison effects of S3I-201, Stattic, IgG, anti-IL11, and anti-IL11RA on Caspase3 activation and CHOP induction. (**E-F**) Quantification of (E) $\alpha SMA^{+ve}$ cells and (F) Collagen I immunostaining from TGFβ1- or rhIL11- stimulated HCFs; data are shown as box-and-whisker with median (middle line), 25th–75th percentiles (box) and min-max percentiles (whiskers); one-way ANOVA with Tukey's correction. (A-F) primary HCFs; 24h; IL11/TGFβ1 (10 ng/ml), U0126 (10 μM), S3I-201 (20 μM), Stattic (2.5 μM), IgG/anti-IL11/anti-IL11RA (2 μg/ml), Tunicamycin (5 μg/ml), Thapsigargin (300 nM) unless otherwise specified. BL: baseline

IL11 (mouse)
Proline content = 8.54%

```
            1          11         21         31         41         51
            |          |          |          |          |          |
  1 MNCVCRLVLV VLSLWPDRVV APGPPAGSPR VSSDPRADLD SAVLLTRSLL ADTRQLAAQM    60
 61 RDKFPADGDH SLDSLPTLAM SAGTLGSLQL PGVLTRLRVD LMSYLRHVQW LRRAGGPSLK   120
121 TLEPELGALQ ARLERLLRRL QLLMSRLALP QAAPDQPVIP LGPPASAWGS IRAAHAILGG   180
181 LHLTLDWAVR GLLLLKTRL
```

**Figure S3. Amino acid sequence of mouse IL11.** Proline residues highlighted in yellow.

**Materials and Methods**

***Antibodies***
BIP (3177, CST), Cleaved-Caspase 3 (9664, CST), Caspase 3 (9662, CST), CHOP (5554, CST), Collagen I (ab34710, Abcam), phospho-ERK1/2 (4370, CST), ERK1/2 (4695, CST), GAPDH (2118, CST), neutralizing anti-human gp130 (MAB628, R&D Systems), anti-gp130 (PA5-99526, Thermo Fisher), gp130 (PA5-28932, Thermo Fisher, IF), IgG (11E10, Aldevron), neutralizing anti-IL11 (X203, Aldevron), commercial anti-IL11 (MAB218, R&D Systems), neutralizing anti-IL11RA (X209, Aldevron), IL11RA (ab125015, abcam, IF), phospho-mTOR (2971, CST), mTOR (2972,CST), phospho-p70S6K (9205, CST), p70S6K (2708, CST), phospho-S6 ribosomal protein (4858, CST), S6 ribosomal protein (2217, CST), $\alpha$-SMA (ab7817, Abcam, Operetta), $\alpha$-SMA (19245, CST, WB), phospho-SMAD2 (3108, CST), SMAD2 (5339, CST), phospho-STAT3 (4113, CST), STAT3 (4904, CST), XBP-1 (sc-8015, SantaCruz), mouse Alexa Fluor 488 secondary antibody (ab150113, Abcam), mouse HRP (7076, CST), rabbit Alexa Fluor 488 secondary antibody (ab150077, Abcam), rabbit HRP (7074, CST).

***Recombinant proteins***
Commercial recombinant proteins: Recombinant human TGFβ1 (PHP143B, Bio-Rad).
Custom recombinant proteins: Recombinant human IL11 (rhIL11, UniProtKB:P20809) and recombinant mouse IL11 (rmIL11, UniProtKB: P47873) were synthesized without the signal peptide. HyperIL-11 (IL11RA:IL-11 fusion protein), was constructed using a fragment of IL11RA (amino acid residues 1–317; UniProtKB: Q14626) and IL-11 (amino acid residues 22–199, UniProtKB: P20809) with a 20 amino acid linker (GPAGQSGGGGGSGGGSGGGSV) [1]. All custom recombinant proteins were synthesized by GenScript using a mammalian expression system.

***Chemicals***
Cycloheximide (C1988, Sigma), 4',6-diamidino-2-phenylindole (DAPI, D1306, Thermo Fisher), Halofuginone (sc-211579, SantaCruz), Nintedanib (S1010, Selleck Chemicals), Pirfenidone (P2116, Sigma), S3I-201 (SML0330, Sigma), Stattic (S7947, Sigma), U0126 (9903, CST), Thapsgargin (12758, CST), Tunicamycin (12819, CST).

### Cell culture
Primary adult human cardiac fibroblasts (HCFs), primary adult mouse cardiac fibroblasts (MCFs), and primary adult human lung fibroblasts (HLFs) were grown and maintained at 37°C and 5% $CO_2$. The growth medium was renewed every 2–3 days and cells were passaged at 80% confluence, using standard trypsinization techniques. All experiments were carried out at low cell passage (<P3). Cells were serum-starved overnight in basal media prior to stimulation. Cells were stimulated with different treatment conditions and durations, as outlined in the main text or

figure legends. Stimulated cells were compared to unstimulated cells that have been grown for the same duration under the same conditions, but without the stimuli.

***Primary HCF culture***

Primary HCFs (6330, ScienCell) were grown and maintained in FM-2 complete media which contains Fibroblast medium-2 (2331, ScienCell), Fibroblasts growth supplement-2 (FGS-2, 2382, ScienCell), 5% fetal bovine serum, and 1% Penicillin-streptomycin (P/S, 0353, ScienCell).

***Primary MCF culture***

Primary MCFs were isolated from *Il11ra1*-deleted mice (*Il11ra*$^{-/-}$) and wild-type (WT, *Il11ra*$^{+/+}$) mice (B6.129S1-*Il11ra*$^{tm1Wehi}$/J, Jackson's Laboratory). Atria were minced and digested with mild agitation for 30 min at 37°C in Dulbecco's Modified Eagle Medium (DMEM, 11995-065, Gibco) containing 1% P/S and 0.14 Wunsch U ml$^{-1}$ Liberase (5401119001, Roche). Fibroblasts were enriched via negative selection with magnetic beads against mouse CD45 (leukocytes), CD31 (endothelial) and CD326 (epithelial) using a QuadroMACS separator (Miltenyi Biotec) according to the manufacturer's protocol. Fibroblasts were maintained and cultured in complete DMEM supplemented with 10% FBS and 1% P/S.

***Primary HLF culture***

Primary HLFs (CC-2512, Lonza) were grown and maintained in FGM-2 complete media, which contains Fibroblast basal medium (CC-3131, Lonza) and hFGF-B, Insulin, fetal bovine serum, GA-1000 as growth supplements (FGM™-2 SingleQuots™, CC-4126, Lonza).

**Half Maximal Inhibitory Concentration Measurement**

HCFs and MCFs were stimulated with IL11 (24 hours) in the presence of IgG (11E10, 2 µg/ml; Aldevron) and varying concentrations of gp130 antibodies (MAB628 for HCFs and PA5-99526 for MCFs; 61 pg/ml to 4 µg/ml; 4-fold dilutions). Supernatants were collected and assayed for the amount of secreted MMP2. Dose-response curves were generated by plotting the logarithm of the gp130 antibodies tested concentrations (pM) versus the corresponding inhibition values (%), using the least squares ordinary fit. We considered MMP2 secretion by unstimulated cells as maximal inhibition (100%), while MMP2 secretion after stimulation with IL11 constituted 0% inhibition.

**siRNA Knockdown**

HCFs were transfected using Lipofectamine RNAiMax (Life Technologies), following the manufacturer's instructions for reverse transfection. Cells were transfected with 12.5 nM On-Targetplus siRNAs (siEPRS: L-008245-00-0005, siNT: D-001810-10-05, Dharmacon) in a medium consisting of serum-free Opti-MEM and complete FM-2 media in a 1:9 ratio. Following 24 hours of transfection, cells were serum-starved overnight prior to stimulation with IL11, HyperIL11 or TGFβ1.

**Operetta high throughput phenotyping assay**

The Operetta assay was performed as previously described [2] with minor modifications: HCFs were seeded in 96-well black CellCarrier plates (PerkinElmer) at a density of $5x10^3$ cells per well. Following simulations, cells were fixed in 4% PFA (Thermo Fisher), permeabilized with 0.1% Triton X-100 (Sigma) and non-specific sites were blocked with 0.5% BSA and 0.1% Tween-20 in PBS. Cells were incubated overnight (4°C) with primary antibodies (1:500), followed by incubation with the appropriate Alexa Fluor 488 secondary antibodies (1:1000). Cells were counterstained with 1 µg/ml DAPI (D1306, Thermo Fisher) in blocking solution. Each condition was imaged from duplicated wells and a minimum of 7 fields/well using Operetta high-content imaging system 1483 (PerkinElmer). Cells expressing ACTA2 were quantified using Harmony v3.5.2 (PerkinElmer) and the percentage of activated fibroblasts/total cell number ($\alpha$-SMA$^{+ve}$) was determined for each field. The measurement of fluorescence intensity per area (normalized to the number of cells) of Collagen I was performed with Columbus 2.9 (PerkinElmer).

**Live/Dead Cells quantification assay**
HCFs were seeded in 96-well black CellCarrier plates at a density of $6x10^3$ cells/well. Following stimulation, cells were incubated 1 h with 1 µg/ml Hoechst 33342 (62249, Thermo Fisher Scientific) and DRAQ7 (D15106, Thermo Fisher Scientific; 1µg/ml) in serum-free basal medium. Each condition was imaged from 3 wells and a minimum of 23 fields/well using Operetta high-content imaging system (1483, PerkinElmer). Live and dead cells were quantified using Harmony v3.5.2 (PerkinElmer).

**Enzyme-linked immunosorbent assay (ELISA)**
The levels of IL11 and MMP2 in equal volumes of cell culture media were quantified using Human IL-11 Quantikine ELISA kit (D1100, R&D Systems) and Total MMP-2 Quantikine ELISA kit (MMP200, R&D Systems), respectively according to manufacturer's instructions.

**Immunoblotting**
For protein extraction and immunoblotting purposes, cells were seeded on 6-well plates at a density of $2x10^5$ cells. HCFs or MCFs were homogenized in RIPA Lysis and Extraction Buffer (89901, Thermo Scientific) containing protease and phosphatase inhibitors (Roche). Protein lysates were then separated by SDS-PAGE, transferred to PVDF membranes, blocked for 1 hour with 3% BSA, and incubated overnight with the primary antibodies (1:1000). Protein bands were visualized using the ECL detection system (Pierce) with the appropriate HRP (1:5000).

**Quantitative polymerase chain reaction (qPCR)**
Total RNA was extracted from HCF lysate using RNeasy column (Qiagen) purification and cDNAs were synthesized with iScript™ cDNA synthesis kit (Bio-Rad) according to manufacturer's instructions. Gene expression analysis was performed with fast SYBR green (Qiagen) technology using StepOnePlus™ (Applied Biosystem) over 40 cycles. Expression data were normalized to *GAPDH* mRNA expression and

fold change was calculated using $2^{-\Delta\Delta Ct}$ method. The primer sequences are as follow:

| Gene | Forward | Reverse |
|---|---|---|
| *EPRS* | 5'-GTGTTTGGGCCACCCTAAAAG-3' | 5'-CTGGAGGAAATCTGACGGTAAC-3' |
| *GAPDH* | 5'-GGAGTCAACGGATTTGGTCG-3' | 5'-ATCGCCCCACTTGATTTTGG3' |

**OP-Puro Protein Synthesis Assay**
HCFs were seeded in 96-well black CellCarrier plates (PerkinElmer) at a density of $5x10^3$ cells per well. Cycloheximide (100 µg/ml), a translation inhibitor used as a negative control, was added to the cells in basal medium, one hour prior to OPP labelling. OPP labelling was performed during the last treatment hour using Click-iT® OPP reagent (C10456 kit, ThermoFisher Scientific) at a final concentration of 20µM in basal medium. After incubation, cells were washed once with PBS and fixed using 4% paraformaldehyde in PBS for 15min at room temperature (RT), and permeabilized using 0.5% Triton® X-100 for 15min at RT. 100µl Click-iT® OPP reaction cocktail containing Alexa Fluor™ 488 picolyl azide was added into each well and incubated for 30 mins at RT, protected from light. Cells were washed with Click-iT® Reaction Rinse Buffer before proceeding for DNA staining, using HCS NuclearMask™ (1µg/ml) Blue Stain following standard procedures. Images were acquired using the Operetta High Content Imaging system.

**Statistical Analyses**
Statistical analyses were performed using GraphPad Prism software (version 8). Statistical significance between control and experimental groups were analysed by two-sided Student's t tests or by one-way ANOVA as indicated in the figure legends. P values were corrected for multiple testing according to Dunnett's (when several experimental groups were compared to a single control group) or Tukey (when several conditions were compared to each other within one experiment). The criterion for statistical significance was $P < 0.05$.


1. Dams-Kozlowska H, Gryska K, Kwiatkowska-Borowczyk E, Izycki D, Rose-John S, Mackiewicz A (2012) A designer hyper interleukin 11 (H11) is a biologically active cytokine. *BMC Biotechnol* **12**: 8.
2. Schafer S, Viswanathan S, Widjaja AA, Lim W-W, Moreno-Moral A, DeLaughter DM, Ng B, Patone G, Chow K, Khin E, et al. (2017) IL-11 is a crucial determinant of cardiovascular fibrosis. *Nature* **552**: 110–115.
